# A prebiotic-postbiotic combination supports dietary carbohydrate-targeting functional properties in a fiber-deprived microbiota

**DOI:** 10.1101/2025.09.23.677878

**Authors:** Amrei Rolof, Erica T. Grant, Oskar Hickl, Stéphanie Willieme, Alessandro de Sciscio, Alina Burghard, Clara Delaroque, Alex Steimle, Richard Ammer, Uwe Baumann, Mahesh S. Desai

**Author notes:** **Correspondence:** (M.S. Desai).

## Abstract

Dietary fiber deprivation compromises gut mucosal barrier integrity by promoting microbial degradation of host mucus, a process linked to various gut-related auto immune diseases. While postbiotics are considered safer alternatives to fiber for susceptible patients, their mechanistic effects on a fiber-deprived gut remain poorly understood. Here, we demonstrate in a mouse model that a fermented postbiotic, alone or in combination with a prebiotic and aloe vera, counteracted the increase of detrimental properties of the microbiota on a fiber-free diet. The supplement regimen reshaped the gut microbiota, counteracting the expansion of key mucin-degrading bacteria, including *Akkermansia muciniphila* and *Parabacteroides goldsteinii*. Metatranscriptomic analysis revealed this compositional change corresponded to a community-wide functional pivot away from expressing mucinolytic enzymes, such as sialidases, and towards utilizing alternative substrates. These microbial shifts recapitulated the effects of dietary fiber reintroduction and translated to direct host benefits, including sustentation of the colonic mucus layer and attenuation of diet-induced type III immune cytokine expression. Our findings provide a mechanistic rationale for using postbiotics to functionally replace dietary fiber, offering a promising strategy to support gut homeostasis in contexts where fiber intake is limited.

## Introduction

Gut microbiome alterations and intestinal barrier dysfunction are associated with various autoimmune, inflammatory and metabolic diseases, including irritable bowel syndrome (IBS), inflammatory bowel diseases (IBD), multiple sclerosis, rheumatoid arthritis, systemic lupus erythematosus, and type 1 diabetes [1–8], with direct causal evidence provided by spontaneous colitis development in Muc2-deficient mice, the dominant mucin type in the lower gastrointestinal tract [9]. The intestinal barrier—composed generally of the epithelial layer and an overlaying mucus layer—exhibits regional differences in thickness and glycan composition along the gastrointestinal tract [10]. In the small intestine, a thin, loosely attached mucus layer facilitates nutrient absorption, while in the colon, a thicker mucus layer physically separates host epithelium from luminal microbes [10]. The mucus forms a hydrogel, mainly composed of mucin glycoproteins together with micro- and macromolecules, such as salts and proteins. This complex network creates a supportive habitat of a dense microbial community under slow-transit conditions [10]. The detrimental effects of reduced barrier integrity have been extensively studied in preclinical mouse models of colitis, which is driven by diet-induced alterations in the gut microbiota metabolic activities [11–15]. Hence, modulating the microbiota composition and activity through dietary strategies represents a promising approach to support mucosal barrier function [16]. Such strategies could include the incorporation of dietary supplements like prebiotics [17, 18] or postbiotics [19, 20], as well as other potentially microbiome modulating supplements that have been shown to improve gut barrier function, such as aloe polysaccharides [21, 22]. While microbiome modulation may have beneficial implications in various diseases, our particular focus lies on IBS, one of the most common gastrointestinal disorders with a prevalence of 4.1 % [23]. Currently, no universal treatment exists due to its multifactorial etiology, highlighting the need for intervention strategies supporting gut barrier integrity.

Fermented foods such as yoghurt, kefir, kimchi, sauerkraut, sourdough, and kombucha result from fermentation and have been part of the human diet for millennia [24–26]. Removal of fermentable substrates and removal or inactivation of life bacteria yields a "postbiotic", which can still contain inactive bacteria or cell wall components [17, 27]. Fermentation generates bioactive compounds, that can render potential health benefits—such as short-chain fatty acids (SCFAs) and transformed polyphenols [18, 28–34]. Most studies to date have emphasized on the effect of non-pasteurized probiotic foods on the gut microbiome [35–39], which may be well-tolerated in healthy populations, but can harbor risks in susceptible individuals [17, 40]. Therefore, recent scientific interest has risen with a focus on interactions between postbiotics with a higher safety profile, the host, and the gut microbiome [19, 20, 41].

Some microbial metabolites, like SCFAs, directly interact with host intestinal G-protein coupled receptors, including GPR43. This can indirectly modulate intestinal barrier function by the promotion of regulatory T cell differentiation, leading to enhanced mucus production by goblet cells and attenuated transcellular permeability [42–44]. Moreover, SCFAs contribute to an acidic luminal environment, thereby supporting the gel-like properties of mucus, improving the physical gut barrier function [45]. Alongside these direct effects on the host, bioactive molecules transiting the gastrointestinal tract can also interact with resident gut microbes, leading to shifts in microbial composition. Explicitly, this has been demonstrated through supplementation of purified bioactive molecules such as SCFAs [20, 46–49], resveratrol [50], secondary bile acids [51, 52], and polyamines [53, 54]. Additionally, several polyphenols show microbiome modulating effects in *in vitro* fermentation systems [55], and a systematic review suggests similar effects by polyphenol-rich food in humans [56]. Regular consumption of fermented foods has been found to increase the abundance of butyrogenic genera from the

*Lachnospiraceae* family[39], including *Anaerostipes*, *Butyricimonas*, *Lactiplantibacillus* and *Eubacterium* [57, 58]. Simultaneously, it also leads to a decrease in potentially pro-inflammatory taxa such as members of the *Clostridium* and *Lachnoclostridium* genus [57–59], which are typically enriched in obese individuals [60] and those consuming a Western-style diet [61, 62]. However, specific mechanistic impacts, particularly for non-SCFA metabolites, remain unclear [17, 18, 36, 63, 64]. Furthermore, little is known, either in mice or in humans, about how postbiotics shape the microbiome under modern, Western-style dietary conditions. Existing research focused mainly on heat-inactivated bacteria or purified bacterial protein as postbiotics [65, 66]. However, this dietary component is of high relevance considering reported low fiber intake among patients suffering from bowel problems [67, 68]. Here, we test different combinations of a fermented, postbiotic (FP) alone, or in combination with only a prebiotic product (PP), or both additionally with an aloe vera juice (AV), in addition to various other purported health benefits [69–71]. The effects of each single product as well as their combination—together with the effect of dietary factor—on the gut microbiome and host barrier integrity are unknown.

To elucidate specific effects on the gut microbiome composition, we supplemented specific-pathogen-free mice with the FP—alone, with the PP, and with both the PP and AV—while undergoing no, short-term, or long-term fiber-deprivation. By analyzing both longitudinal and cross-sectional data after two weeks of supplementation, we observed specific supplement-induced microbial shifts in fiber-deprived mice resembling those observed in short-time deprived mice after full reintroduction of dietary fiber. These shifts included mitigating the fiber-free diet-induced proliferation of mucin-degrading species such as *Akkermansia muciniphila* and *Parabacteroides goldsteinii*, as well as the increase in certain butyrate-producing species encompassing genera such as *Enterocloster*, *Dysosmobacter* and *Kineothrix*. Furthermore, the abundance shifts were accompanied by a lower predicted fecal abundances in sialidases, specific mucin-targeting carbohydrate-active enzymes (CAZymes), compared to FF-fed control mice. We confirmed this product-mediated reduction in sialidase transcripts in cecal contents, and simultaneously detected increases in several fiber-degradation associated pathways across all supplementation groups. Furthermore, histological analysis of rectal and distal colonic tissues suggested a trend towards a thicker mucus layer of mice receiving all three supplements compared to FF controls with a typically thin mucus layer [11, 13]. This was accompanied by a trend toward reduced expression of several membrane integrity markers and host cytokines in colonic tissues, that are typically connected with a type III immune response. Together, our findings suggest that a postbiotic alone or in combination with a prebiotic and AV juice can modulate functions of pre-perturbed communities by reducing the abundance of mucin-degrading bacteria and enhancing the abundance of SCFA-producing fiber-degrading bacteria. Our data show that targeted microbiota changes by dietary supplements have the potential to support gut homeostasis, therefore providing insights that could help to manage diseases associated with gut barrier impairment.

## Material and Methods

### Specific-pathogen-free (SPF) experiments

The protocols for SPF animal experiments were first pre-approved by the Animal Welfare Service at the Luxembourg Institute of Health (LIH), followed by approval by the Veterinary Services Administration within the Ministry of Agriculture (under the national authorization no. LUPA2024/11). Six-week-old female C57BL/6J mice, purchased from Charles River Laboratories, France, were housed in the LIH SPF facility. The fiber-rich (FR) diet is the SAFE 150 Standard Diet (SAFE S.A.S, Augy, France, E8404A01R 00029). The mice were fed FF diet, which was custom-manufactured by Envigo (Inotiv, Venray, Netherlands), based on the TD.140343 diet formulation, that was previously described [11]. The diets were provided *ad libitum* to each cage together with autoclaved water. Food and water consumption were monitored on a weekly basis. Fecal samples were collected at least on a weekly basis for compositional analysis and up to daily in case of voluntary dropping during handling. According to Figure 1, mice were split into three diet groups, with one group receiving a baseline FR diet and two groups receiving a baseline FF diet. Mice were fed these diets for 3 weeks (wk -3 to 0) prior to intervention. At the start of the supplementation, mice of the third group (FF-fed) were switched back to a FR diet, the other two groups remained on their respective diets. Throughout the intervention time (6 weeks for the FR-fed and 2 weeks for FF-fed groups), mice received a daily intragastric gavage of a 200 µL gavage mix containing the indicated dose of product(s) dissolved in 1x phosphate buffered saline (PBS), or 200 µL 1x PBS solution as a control. All gavage mixes and control solutions were filter-sterilized, aliquoted into light-protective tubes sealed with parafilm, and stored at 4 *^◦^*C one week prior to the begin of the supplementation period. According to Figure 1, within each diet group, mice were allocated into the following four supplementation groups: 1) control or placebo-intervened (PBS), 2) gavage mix containing only the fermented product (FP), 3) same as group 2 plus a prebiotic product (PP) integrated into the diet, and 4) same as group 3 plus aloe vera (AV) in the gavage mix. The PP was integrated into the diet by the manufacturer of each specific diet at a concentration of 2.68 g/kg. The experiment was ended for all FF fed mice due to a stress-related weight loss, that was most likely caused by the combination of daily hand restraining and gavage coupled with the poor diet [72], with no differences between supplemented and placebo-intervened groups. At the end of the experiment, mice were humanely euthanized by cervical dislocation, followed by tissue collection for experimental readouts.

**Figure 1:**
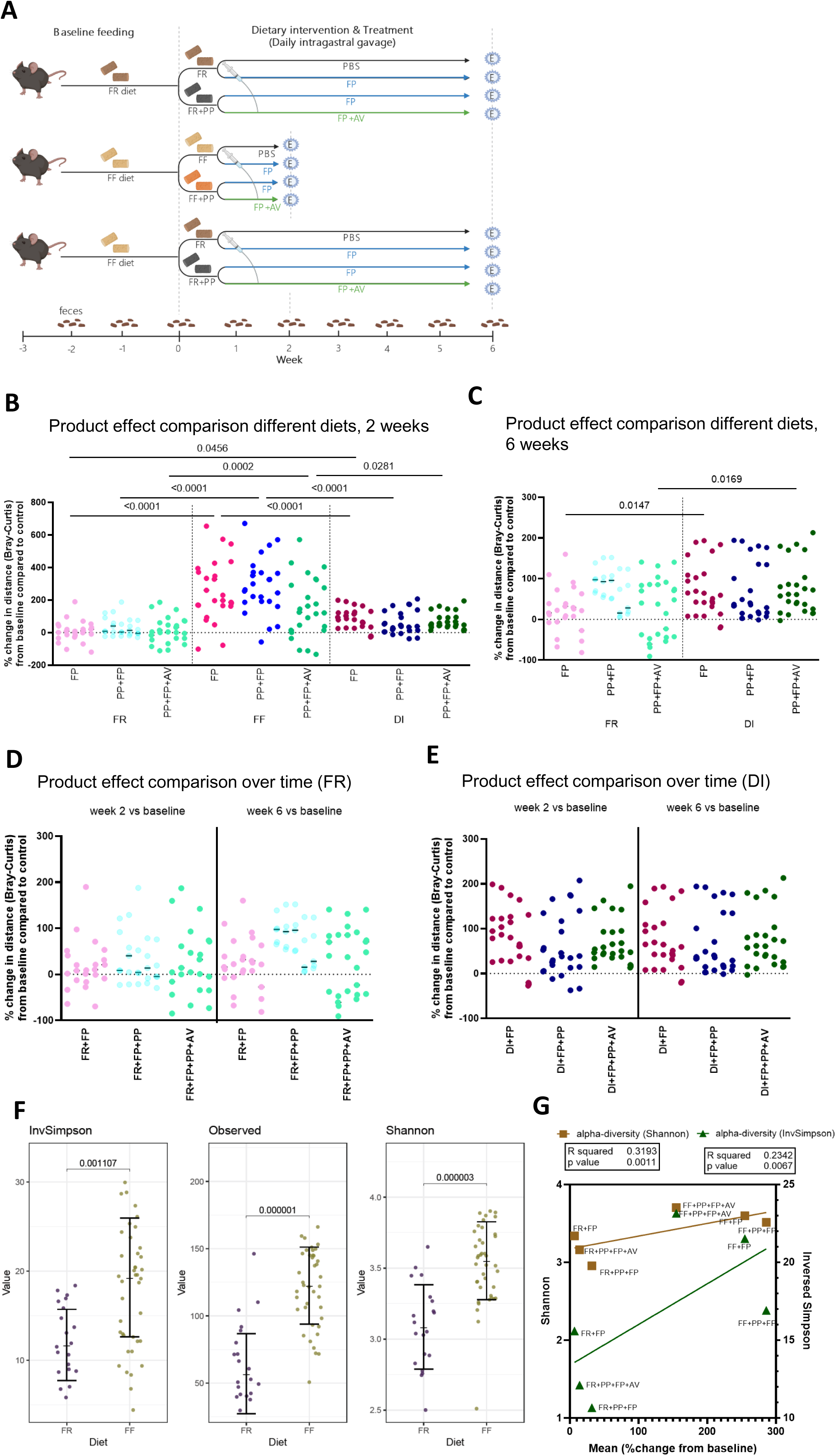
A) Experimental schedule. 6 week-old female specific-pathogen-free (SPF) C57BL/6J mice were baseline fed a fiber-rich (FR) or fiber-free (FF) diet, and either continued on a FR or FF diet, or underwent a dietary intervention (DI) from baseline FF to FR until the end of the experiment. After baseline feeding the supplementation period started, which included the incorporation of a prebiotic product (PP) into the corresponding diet for all PP-supplemented mice, and a daily intra-gastric gavage with a vehicle solution (1x PBS; ctrl group), the fermented product (FP) in 1x PBS solution, or a mix of aloe vera juice (AV) and FP. Supplementation period counted up to 6 weeks in the FR continued groups and 2 weeks in the FF continued group. Fecal samples of all mice were collected at baseline (wk 0), and then weekly during the supplementation. At the end of the supplementation period, mice were sacrificed for various readouts (n = 60). **B-E)** Individual dot plot showing the percent change in distances after B) 2 weeks or C-E) 6 weeks supplementation period of changes in baseline samples in each mouse compared to itself and other individuals in its group (n (individual comparisons)= 5, n (group-intern comparisons) = 20). Measurements are based on the PCoA of the dissimilarity matrix of all groups at wk0 (n=60), wk2 (n=59), wk6 (n=60) (n(total)=179). Statistical analysis of differences within the same supplementation B-C) between diets or D-E) between 2 weeks and 6 weeks within the same dietary background was performed using a two-way ANOVA followed by post hoc multiple comparisons of similar supplements between two diets with false discovery rate (FDR) correction using the Benjamini–Hochberg method (1 family, 9 comparisons, FDR Q = 0.05). **F)** Individual dotplots of alpha- diversity measures (Inverse Simpson, Observed, Shannon) for each mouse at baselines with means and standard error (SE). Statistical analysis was performed using Kruskal–Wallis rank sum test and Dunn’s post-hoc test for pairwise comparisons between diet groups with false discovery rate (FDR) correction using the Benjamini–Hochberg method. Ajusted P.values are indicated between groups. **G)** Average dot plot with simple linear regression of mean alpha-diversity measures (Shannon (left y-axis) and Inversed Simpson (right y-axis) per supplement receiving group, with the same’s group mean percent change from baseline after 2 weeks of supplementation compared to control (x-axis). Calculation performed with 95 % confidence intervall (CI), R^2^ correlation coefficient and t-test on non-zero slope p-value *<*0.05.

### Extraction of genomic DNA and full-length 16S rRNA gene amplification

Immediately after collection, fecal pellets were stored at -20 *^◦^*C, while small intestinal contents were flash-frozen in liquid nitrogen, and stored at -80 *^◦^*C until DNA extraction. DNA extraction from single fecal pellets or total small intestinal contents followed the phenol:chloroform extraction protocol described by Steimle et al. [73]. Genomic DNA purification was performed by using the DNeasy Blood & Tissue Kit (QIAGEN Benelux B.V., Netherlands, Cat. No. / ID: 69506) following manufacturer’s instruction. Extracted dsDNA elution was performed in 50 µL nuclease-free water (Invitrogen™, cat#10977049, FisherScientific, Belgium) and concentration determined with a N60 Micro-Volume UV-VIS Spectrophotometer (Implen™, cat#15442203, FisherScientific, Belgium). Full-length 16S rRNA gene was amplified (Primers: Table 2) following a protocol with an initial 94 *^◦^*C pre-denaturation step for 2 min, followed by 35 cycles of (1) 30 seconds denaturation at 94 *^◦^*C, (2) 30 seconds primer annealing at 51 *^◦^*C, and (3) 90 seconds extension time at 72 *^◦^*C. Amplification success was verified by DNA gel electrophoresis on a 1.5 % agarose gel containing 1X TAE buffer with 1X SYBR Safe DNA gel stain. The running buffer was 1X TAE, and an Invitrogen™ 1 kb plus DNA ladder served as a size control. 16S rRNA gene amplicons with expected size of approximately 1500 bp were cut from the gel using a scalpel, and purified using the Macherey Nagel NucleoSpin® Gel and PCR Clean-up Kit by following the manufacturer’s manual. Amplicon dsDNA was eluted in 20 µL of the manufacturer’s elution buffer (NE) and concentrations determined using a Qubit® 4.0 fluorometer (Invitrogen, cat#33226, FisherScientific, Belgium) .

### Full-length 16S rRNA gene sequencing, sample preparation and data processing

Full-length 16S rRNA gene sequencing of a ^~^50 fmol concentrated library of ultra II end-prepped (NEB, UK) amplicons was performed using the Native Barcoding Kit 96 V14 (Oxford Nanopore Technologies, UK, (SQK-NBD114.96)), following the supplier’s manuals with the use of the Short Fragment Buffer (SFB) and a Flow Cell R10.4.1 (FLO-MIN114, Oxford Nanopore Technologies, UK) on a MinION sequencing device (Oxford Nanopore Technologies, UK) in 400 bps output mode. Raw sequencing data in POD5 format were processed using Dorado v0.9.1 [74].

Basecalling was performed with the ’*dna_r_*10.4.1*_e_*8.2_4_00*bps_s_upv*5.0.0‘ model, and reads were de- multiplexed with the ‘SQK-NBD114-96‘ kit. During this step, reads with a mean quality score below 7 were discarded, and adapter sequences were trimmed. Following initial demultiplexing, unclassified reads were subjected to a second demultiplexing round to recover additional barcoded reads. The resulting FASTQ files for each sample were then processed for quality control using Chopper v0.9.1 [**?**]. Reads with an average quality score below 10 were removed. Furthermore, reads were filtered by length to retain only those between 700 bp and 1900 bp for downstream analysis. For samples processed across multiple sequencing runs, the filtered FASTQ files corresponding to the same sample were concatenated prior to taxonomic analysis.

### Taxonomic Profiling

Taxonomic classification of the processed long reads was conducted using Emu v3.5.1 [75] using a custom reference database based on the GreenGenes2 database (v2024.09) [76]. The database was built using the full-length 16S rRNA gene sequences and the corresponding taxonomy provided by GreenGenes2. Taxon abundance for each sample was quantified against the GreenGenes2 database using the ‘emu abundance‘ command with the ‘–type map-ont‘ setting.

### Functional Profile Prediction

The functional potential of microbial communities was inferred from the 16S rRNA gene data using PICRUSt2 v2.6.2 [77]. The taxonomic profiles generated by Emu were processed to create the required abundance table and sequence file inputs. The core PICRUSt2 pipeline (‘*picrust*2*_p_ipeline.py*‘) was then executed with this study-specific abundance table and sequence file. The analysis was run in stratified mode (‘–stratified‘) to obtain per-taxon contributions to the overall functional profiles. Predictions were made for multiple gene and pathway databases, including the Kyoto Encyclopedia of Genes and Genomes (KEGG) Orthology (KO), Enzyme Commission (EC) numbers, Gene Ontology (GO) terms, Pfam, BiGG, and CAZy [78–81]. Finally, key PICRUSt2 output files, including the predicted metagenome contributions and NSTI (Nearest Sequenced Taxon Index) values, were annotated with the full taxonomic lineages from the GreenGenes2 database to facilitate interpretation.

### RNA extraction and reverse transcription

Following dissection, large intestinal tissue samples were placed in RNAprotect Tissue Reagent (QIAGEN, cat# 76104, Netherlands), while cecal contents were mixed with RNAprotect Bacteria Reagent (QIAGEN, cat# 76506, Netherlands) and incubated at 4 *^◦^*C overnight. After removing the supernatant, all samples were stored at -80 *^◦^*C until processing. Cecal contents were thawed on ice and combined with Type A Ceramic Beads (0.6–0.8 mm, Macherey Nagel, Cat# 740786.B.250), 500 µL Buffer A (200 mM NaCl, 200 mM Tris, 20 mM EDTA), 210 µL SDS (20 % w/v, filter-sterilized), and 500 µL phenol:chloroform:isoamyl alcohol, pH 4.3 (Carl Roth, Cat# 985.2). Homogenization was performed using a RETSCH Mixer Mill MM 400 for 3 minutes at room temperature, followed by centrifugation at 18,000 × g for 3 minutes at 4 *^◦^*C. The aqueous layer was transferred into to a second tube and subjected with an equal volume of phenol:chloroform:isoamyl alcohol, followed by centrifugation under the same conditions. RNA was precipitated by adding one volume of ice-cold 100 % ethanol and one-tenth volume of 3 M sodium acetate, and samples were stored at -80 *^◦^*C for 20 minutes. After centrifugation (18,000 × g, 4 *^◦^*C, 20 min), the supernatant was discarded, and RNA pellets were washed twice with 500 µL of chilled 70 % ethanol, air-dried, and dissolved in 50 µL of nuclease-free water. Samples were incubated at 56 *^◦^*C for 15 minutes to ensure complete dissolution. For colonic tissues, RNA was extracted using Trizol Reagent (ThermoFisher Scientific, Cat# 15596018) with mechanical disruption by metal beads, following the manufacturer’s instructions. RNA concentrations were measured using a NanoPhotometer N60 (Implen). Cecal RNA samples were adjusted to 400 ng/µL, and colonic RNA to 200 ng/µL. Reverse transcription was performed using Super- Script™ IV Reverse Transcriptase (Invitrogen™, cat# 18090200) with 500 µM dNTPs, 2.5 µM random primers, and 40 U RNaseOUT™ (all from Invitrogen™, FisherScientific, Belgium) according to the supplier’s protocol. Finally, complementary DNA was diluted 1:5 in nuclease-free water for downstream qPCR analyses.

### RNA sequencing and analyses

Cecal RNA samples were processed to be sent for sequencing at the LuxGen Platform (Dudelange, Luxembourg). Libraries were prepared using the Illumina Stranded Total RNA Prep with Ribo-Zero Plus kit following the manufacturer’s protocol (Illumina Document 1000000124514 v02, April 2021). Library quality and concentration were evaluated on a Fragment Analyzer using the HS NGS kit, normalized to 10 nM, pooled, and sequenced on an Illumina NovaSeq 6000 SP200 flow cell (paired-end, 75 bp). Raw reads (24–35 million per sample; median 28 million) were filtered using KneadData v0.10.0 to remove low-quality reads and sequences mapping to rRNA or the mouse genome (BowTie2 v1.3.199). Cleaned datasets (5.1–14 million reads; median 9.7 million) were processed in MetaPhlAn 4.1.1 [82] for taxonomic profiling using the "vJun23_CHOCOPhlAnSGB_202403" database. Functional annotation and assignment of gene families to pathways, including stratification by predicted bacterial contributor, were performed in HUMAnN 3.9 [83] with ChocoPhlAn "201901_v31" and UniRef201901b databases. Gene family abundances were converted to counts per million, and UniRef90 identifiers were mapped to CAZy and Pfam categories. Pathway data were retained with default MetaCyc identifiers.

### Quantitative PCR

PCR reactions were prepared in a total volume of 12.5 µL per well, containing 1x PCR Buffer (with MgCl_2_), 2.5 mM MgCl_2_, 400 µM dNTPs (Invitrogen™, Cat# 10297018, FisherScientific, Belgium), 0.2 µM of each forward and reverse primer (Table 2), 1x SYBR Green I Nucleic Acid Gel Stain (Invitrogen™, Cat# S7563, FisherScientific, Belgium), 0.5 U Platinum™ Taq DNA Polymerase (Invitrogen™, Cat# 10966034, FisherScientific, Belgium), and 1 µL of cDNA template (200 ng/µL). Reactions were set up in white PCR plates (Bioplastic, Cat# AB70659) and briefly centrifuged. Amplification was performed on a C1000 Thermal Cycler (Biorad, Cat# 1855195) controlled via Biorad CFX Maestro software (Version 5.3.022 1030). The program included an initial denaturation step at 94 *^◦^*C for 5 min, followed by 40 cycles of 94 *^◦^*C for 20 s, 60 *^◦^*C for 20 s, and 72 *^◦^*C for 45 s, ending with a final extension at 72 *^◦^*C for 5 min. A melting curve was generated by increasing the temperature from 65 *^◦^*C to 95 *^◦^*C in 0.3 *^◦^*C increments, with readings taken every 15 s. Standard curves were prepared by pooling equal amounts of five representative samples (standard 1, 200 ng/µL), followed by three serial 1:2 dilutions (standards 2–4) and four serial 1:4 dilutions (standards 5–8). All standards were run in duplicate alongside the test samples. Pre-analysis was performed to determine Ct-thresholds: dilutions of 1:2 produced an approximate decrease of 1 Ct unit, while 1:4 dilutions produced a decrease of 2 Ct units. For delta Ct calculations, measurements above the Ct-thresholds (Table 2) or values with standard deviations greater than 1 between replicates were excluded. Ct values were corrected for amplification efficiency (E) for both housekeeping and target genes before calculating delta Ct values.

**Table 1:** Per mouse gavage volumes for each experimental group.

| # | Groups | PBS [μL] | FP [μL] | AV [μL] |
| --- | --- | --- | --- | --- |
| 1 | PBS (control) | 200 | 0 | 0 |
| 2 | FP \| FP+PP | 129.6 | 70.4 | 0 |
| 3 | FP+PP+AV | 41.7 | 70.4 | 87.9 |

**Table 2:**
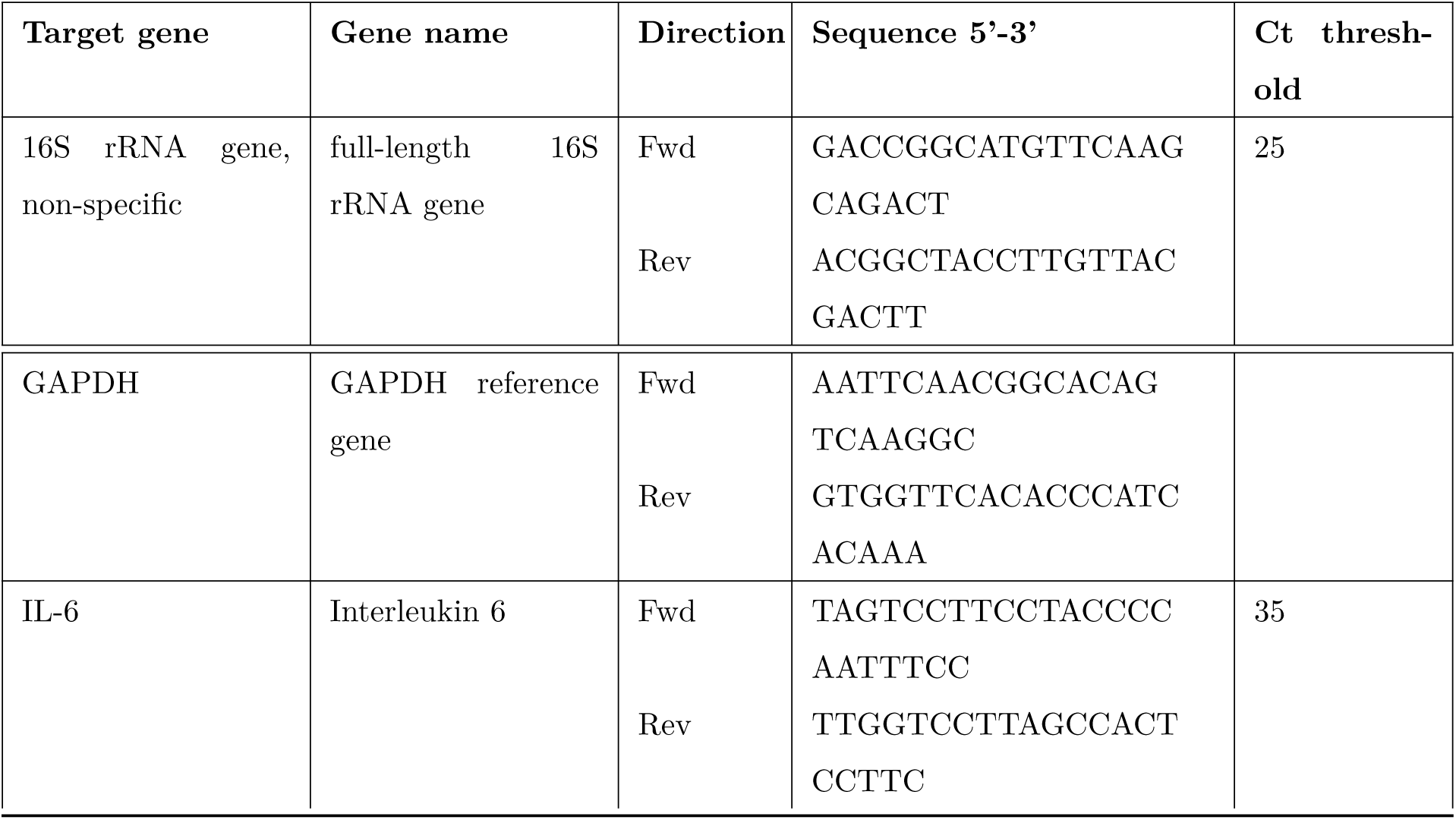
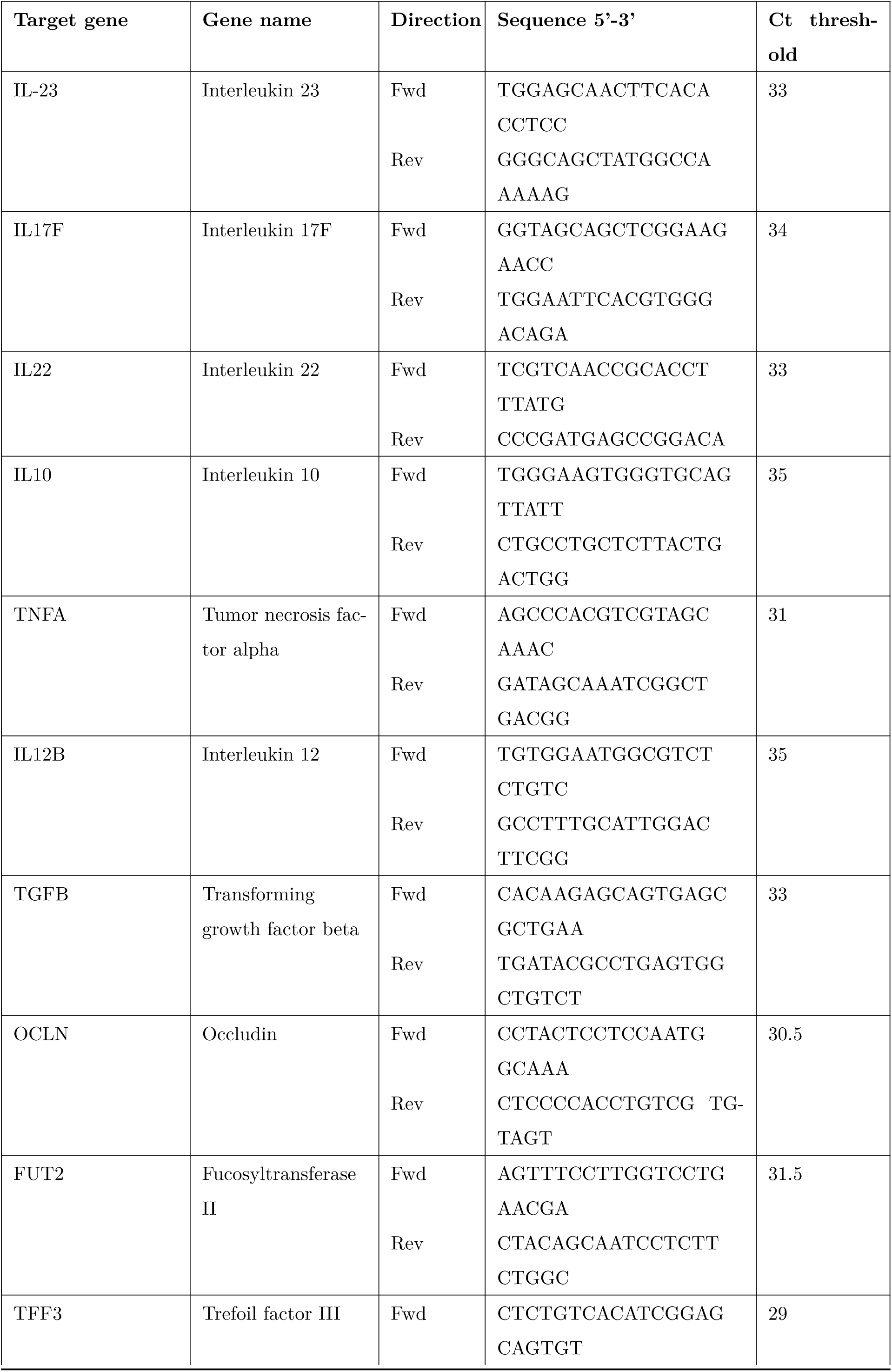
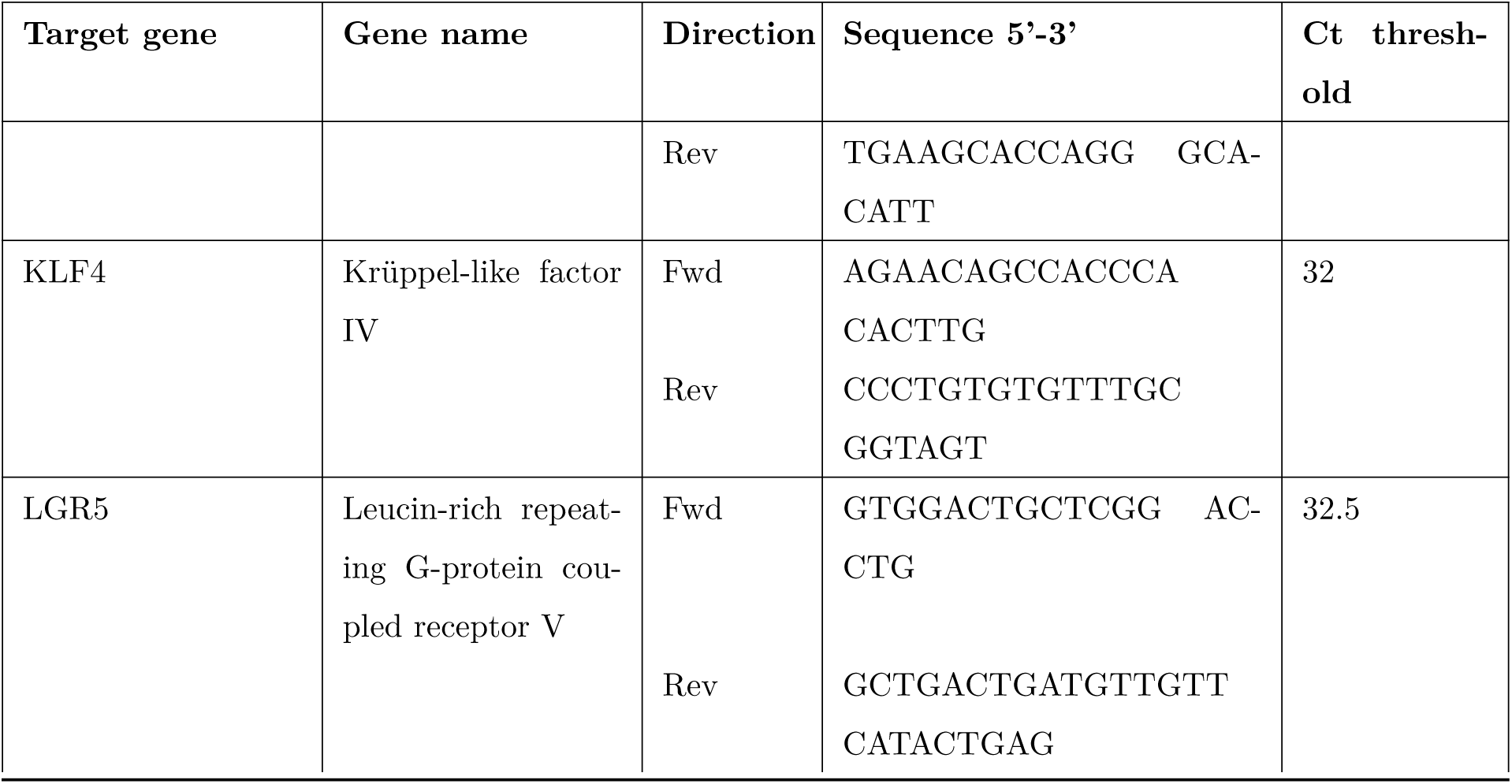
Primer sequences used for PCR and qPCR amplification.

### Histology and Mucus Layer Quantification

Distal colon segments (1–2 cm) were fixed in freshly prepared methacarn solution (methanol:chloroform:acetic acid, 60:30:10 v/v) for 3–5 hours. After fixation, tissues were transferred to 100% methanol for overnight incubation and stored at 4 *^◦^*C in fresh methanol until further processing, as described previously [12]. Samples were embedded in paraffin using a Thermo Excelsior AS Tissue Processor. The automated dehydration and infiltration sequence included three short methanol washes (1 min each), three ethanol washes (1 h each) at 4 *^◦^*C, a single 1-hour xylene wash at 4 *^◦^*C, followed by three 30-minute xylene washes at room temperature and three 1-hour paraffin infiltration steps. Blocks were cut into 4 µm sections on a microtome and mounted on glass slides. Two slides per sample were stained with Alcian Blue (pH 2.5) using the Thermo Gemini AS system. The staining procedure involved two 5-minute washes in Tissue Clear/xylene substitute, sequential ethanol washes (100%, 95%, and 70%; 2 min each), a 3-minute rinse in distilled water, and Alcian Blue staining for 5 min. Slides were rinsed again in distilled water (3 min), counterstained with 0.1% nuclear fast red–aluminum sulfate for 10 min, and subjected to graded ethanol dehydration (70%, 95%, and 100%, twice, 1 min each) before clearing in Tissue Clear/xylene substitute for 5 min. Slides were scanned using the NanoZoomer 2.0 HT at 20× magnification, with focus adjusted manually for each sample. Mucus layer thickness was quantified in intact regions using the QuPath Annotation Tool (QuPath 0.6.0). Reported values represent the mean of 3–38 measurements per section.

### Measurement of Carbohydrate-Active Enzyme and Sulfatase Activities

Enzymatic activities were assessed following the protocol of Steimle-Grant et al. [84], with minor modifications. Fecal samples were first weighed and suspended in 260 µL of detection buffer. After centrifugation at 50 *×* g for 5 minutes at 4 *^◦^*C, 10 µL of the supernatant were collected and stored at -20 *^◦^*C for bacterial count. To the remaining 250 µL of sample, an equal volume of 2× concentrated lysis buffer containing enzyme mix and protease inhibitors was added, yielding the final lysate composition as described previously [84]. Modifications included performing all dilutions and measurements in lysis buffer, except for the substrate solution, which was diluted in detection buffer according to the original method. Protein concentrations that were below optimal levels (applies for sulfatase) were adjusted to a minimum of 15 mg/mL. Samples exceeding this threshold were excluded from sulfatase analysis (FR ctrl, n=4; FF ctrl, n=3; FF FP, n=2; FF PP FP, n=2; FF PP FP AV, n=4), and one sample from the FF PP+FP+AV group was omitted from the entire experiment due to limited fecal material. This approach allowed normalization of enzyme activities both to total protein and bacterial counts per gram of stool.

### Bacterial quantification

For the quantification of bacterial cells, 5 µL of the 10 µL fecal suspension in CAZyme detection buffer (prepared as above) were first diluted 1:100 in 1x PBS to a total volume of 100 µL, and then diluted 1:1000 in 1x PBS with 0.1 x SYBR Green a final volume of 100 µL. 1 µL of 5 different samples were mixed in a similar way but finally diluted in 1 x PBS for the unstained control. Cells were counted using an Agilent NovoCyte Quanteon device with a flow rate of 14 µL/min until 25 µL of the sample were recorded. Gating strategy was determined based on the unstained control. Bacterial count per mL was calculated under consideration of dilution factors.

### Statistical analysis

Statistical analysis was performed with the built-in feature "generatetaxatrendtestlong" for longitudinal compositional data of fecal samples and "generatetaxatestsingle" for cross-sectional compositional data of fecal samples, small intestinal and cecal contents, of the "Microbiome- Stat" R package [85]. This included linear regression on CLR-transformed abundances, with post-hoc multi comparison correction for false-discovery rate by Benjamini-Hochberg with an adjusted value *<*0.05 considered as statistically significant, if not stated otherwise. Statistical analysis of experimental CAZyme assay data and qPCR based delta Ct-values was performed using the Prism GraphPad Software (San Diego, CA, USA). If not stated differently, data was tested for normal distributed and then subjected to a two-way ANOVA with post-hoc correction for multiple comparisons by Benjamini-Hochberg (p*<*0.05 considered statistically significant). Non-normally distributed data was tested using the non-parametric Kruskal-Wallis test with multiple comparison corrections by Benjamini-Hochberg, again adjusted p values *<*0.05 were considered as statistically significant.

## Results and discussion

### Low-fiber perturbed communities more susceptible to modulation by postbiotic-based supplementations

Six-week-old, specific-pathogen-free (SPF) C57BL/6J mice (n=60) were fed either a fiber-rich (FR) diet (n=20) or fiber-free (FF) diet (n=40) for three weeks prior to the start of the intervention (Figure 1A). The FR-fed group (n=20) and one FF-fed group (n=20) remained on their baseline diet during the subsequent intervention period, while the other FF-fed group underwent a dietary intervention (DI)—that is, they were switched from the FF diet to an FR diet—upon starting the intervention (n=20) (Figure 1A). Within each diet arm, mice were split into 4 groups (n=5 for each group) receiving the following interventions daily: 1) control ("Ctrl"), which was placebo-intervened by intragastric gavage with PBS (n=5), 2) fermented product ("FP") by intragastric gavage, 3) same as group 2 with addition of a prebiotic product integrated into the diet ("FP+PP"), or 4) same as group 3 with the addition of aloe vera in the gavage mix ("FP+PP+AV"). The intervention period for mice remaining on the FF diet amounted to two weeks, for all other groups to six weeks (Figure 1A). Fecal samples were collected weekly for microbiome composition analysis.

After two weeks of daily administration, fecal microbiome compositions changed in all diet groups (Supp 1A-C), with a stronger shift between time points for FF and DI groups (Supp 1B- C) compared to FR (Supp 1A). From the microbiome composition changes observed in the FR control mice, we concluded underlying within-group stochastic variation. We therefore present these shifts as Bray-Curtis distances relative to the baseline of the supplemented mice compared to their placebo-intervened controls (Figure 1B). Comparing supplementation effects between all three diets, we observed the strongest shifts in microbial communities in mice maintained on FF diet throughout supplementation (Figure 1B). This indicates that the microbiome of these mice might have been more susceptible to modifications by external factors, compared with mice receiving a FR diet [11, 13, 14], or only previously short-term FF fed mice. Moreover, we observed that FP- and FP-PP-AV-supplemented DI mice showed stronger compositional changes to baseline after 6 weeks compared to similarly supplemented FR-fed mice, as confirmed by two-way ANOVA on the percent change in Bray-Curtis distance from baseline compared to the control, with Benjamini-Hochberg FDR correction for multiple comparisons (p*<*0.05) (Figure 1C) [86]. This suggests that fiber-deprived mice, even after returning back to a FR diet, still show a higher instability in the face of external perturbations. However, there were no additional shifts from the baseline after 6 weeks of supplementation compared to 2 weeks of supplementation, neither in the FR fed mice (Figure 1D, Supp 1D), nor the DI mice (Figure 1E, Supp 1E). Our findings therefore suggest that the gut microbiota of FR-fed mice are overall less susceptible to external perturbations, and that even short-time fiber-deprivation can already lead to an increased susceptibility to external perturbations (Figure 1C).

Depending on the substrate specificity and level of sharing among microbiome members, a FR diet can result in lower alpha-diversity measures than an FF diet [87], as seen here for the FF baseline (FF and DI baseline data combined) concerning evenness and richness, measured as Shannon, inverse Simpson and observed indices (Figure 1F, Supp 1G-I). Correlating the potential to external microbiome modulation with alpha-diversity measures revealed a weak (R2 (Shannon) = 0.3193, R2 = 0.2342), but significant correlation between high diversity in FF-fed mice and greater shifts in microbial composition (Figure 1G). These findings suggest a lower microbiome resilience to external modifiers, a finding aligning with recent evidence. However, the interpretation of high alpha-diversity as a direct and universal indicator of a ’healthy’ microbiome is being re-evaluated[88]. Klimenko et al. reported a positive correlation between baseline alpha-diversity and responsiveness to dietary interventions, which was mediated by a lower average gene count per organism (AGN), and which they identified as an indicator for reduced community resilience [89]. In line with these and our findings, communities of mice previously fed a low-fiber, Western-style diet failed to recover from antibiotic treatment, compared to antibiotic treated mice fed a FR diet [90]. Tanes et al. observed humans with a vegan life style to have much faster recovery following microbiome purge than those on a fiber-free exclusive enteral nutrition (EEN)diet [91]. Overall, these findings strongly suggest that mice fed a baseline fiber-free diet are more susceptible to external perturbations.

### Postbiotic-based supplementation leads to specific microbial changes similar to dietary fiber intervention

After identifying general microbial changes within each diet, we further explored supplementation-dependent shifts in relative abundance of single microbial taxa over time, shown as the average change from baseline for 68 species that differed from the FR control group changes compared to baseline (p<0.1) (Figure 2A). After two weeks of supplementation, linear models for differential abundance (LinDA) [85, 92] of cross-sectional data revealed several trends (p*<*0.1) and significantly differentially abundant taxa (FDR adjusted for multiple comparison by Benjamini-Hochberg p-value*<*0.05, using the built-in "*generate_t_axa_t_est_s_ingle*" function of the "MicrobiomeStat" package) within each dietary background compared to the placebo-intervened control (Figure 2B). We saw a total of 134 trends and additional 38 significant shifts across all dietary backgrounds (Figure 2B). Among those, the majority of shifts were observed for the continuous FF background, with 55 shifts in the FF groups and 56 shifts in the DI groups (Figure 2B). For the FR groups, we observed 37 shifts, with none remaining significant after FDR correction (Figure 2B). To identify taxa whose abundance was consistently modulated by the supplements, we implemented a stringent multi-step filtering strategy on longitudinal data from weeks 0, 1, and 2. This process involved 1) selecting taxa that exhibited a clear temporal trend in at least one supplementation group (absolute coefficient *<*0.5; p*<*0.1); 2) retaining from this subset only those taxa that showed a consistent direction of change across all related supplement groups within a given diet (for a robust FP-effect, an increase in the FP group must also be an increase in the FP+PP and FP+PP+AV groups; for a robust PP-effect an increase must be detected in the PP- and PP+AV-supplemented groups), and 3) we excluded any effect whose coefficient’s error bar overlapped with zero (Figure 2C, Supp 2B-C). This approach was designed to isolate only the most robust changes. To validate these findings, we applied the same filtering and plotting strategy to the cross-sectional data at week 2, confirming the same trends for end point relative abundances (Supp 2A). The resulting taxa are presented alongside comparisons between the placebo-intervened diet groups (FF vs. FR and DI vs. FR) to help differentiate supplement-driven effects from underlying diet-associated changes (Figure 2C, Supp 2A). Ad- justed p.values*<*0.05 were additionally marked by a filled dot, indicating statistical significance. Applying these filtering criteria, we identified three bacterial taxa whose abundances were altered by the FF diet and consistently counteracted by the FP supplementation. Specifically, FP supplementation counteracted the FF-driven increase of two known mucin-degrading bacteria, *A. muciniphila* [11, 93] and *P. goldsteinii* [94, 95], and reversed the FF-driven decrease of an unclassified *Enterocloster* species (Figure 2C, Supp2A). The effect on *A. muciniphila* was particularly notable as this species is known to expand under fiber-free conditions, an event often linked to host-detrimental effects [11, 13–15]. Our data first confirmed that reintroducing fiber in the DI group significantly attenuated the increase of this species compared to mice remaining on the FF diet. We then found that FP supplementation alone recapitulated this protective, diet-induced effect (Figure 2C, Supp2A). The addition of PP enhanced this effect, resulting in a statistically significant difference compared to the FF control (p.adj*<*0.05), whereas the further addition of AV conferred no significant change in the FF context (Figure 2C, Supp2A). However, the effects of the supplements were highly dependent on the dietary background. For instance, while AV had no additional effect on *A. muciniphila* in the FF diet, it induced a positive and negative shift in *A. muciniphila* abundance under the FR and DI conditions, respectively. This indicates that all three supplements can modulate the relative abundance of *A. muciniphila*, but the specific outcome is contingent on the host’s dietary fiber intake.

**Figure 2:**
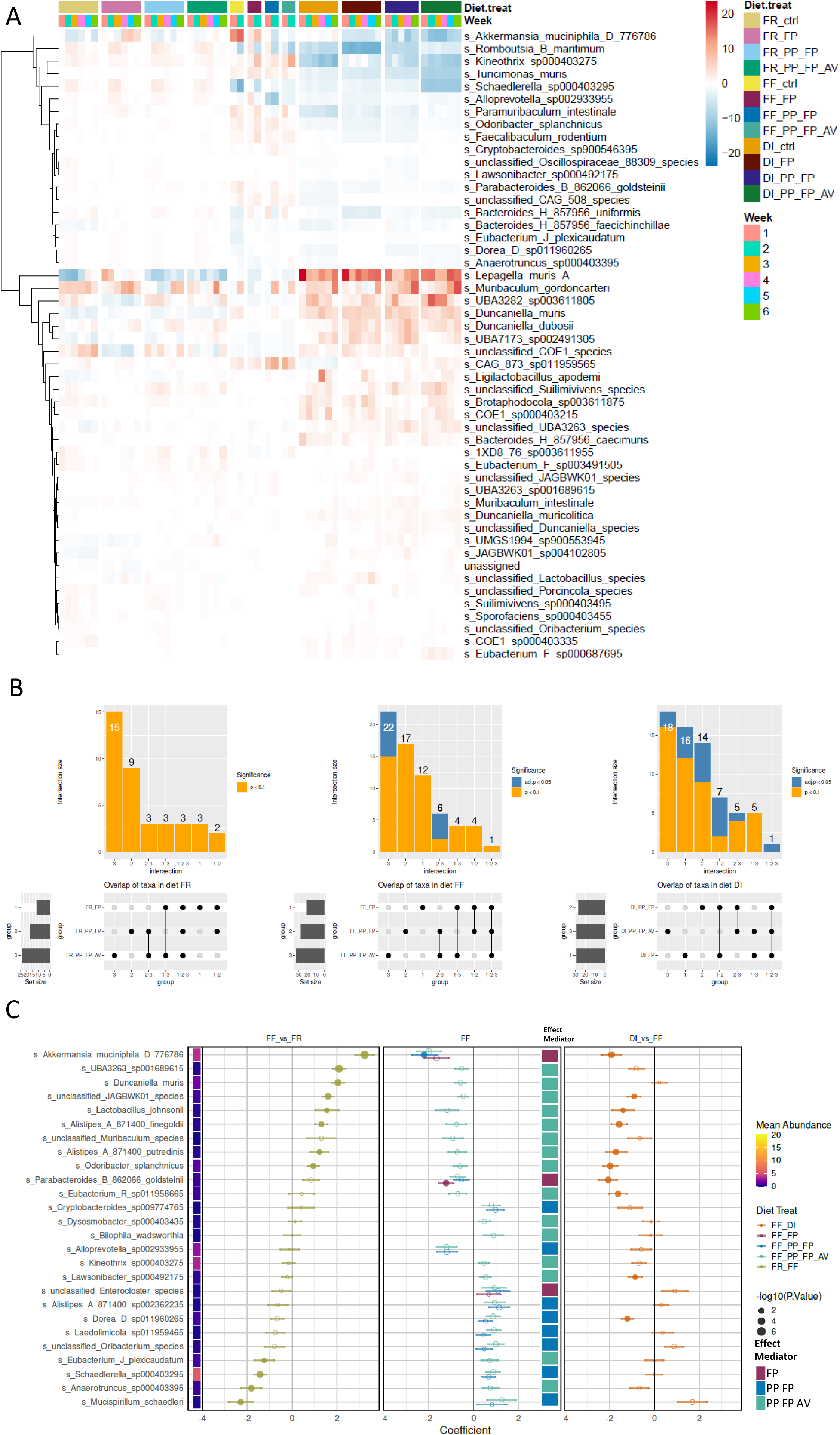
A) Heatmap of mean absolute change in relative abundance (in %) of 50 top abundant species during supplementation period compared to baseline on different dietary backgrounds (fiber-rich (FR), fiber-free (FF) and diet intervention (DI)) and supplementation (fermented postbiotic (FP), prebiotic product (PP), and aloe vera (AV)) vs. PBS control (ctrl) within each diet. **B)** Upset plot showing the number of features with significant change after 2 weeks of supplementation in linear models for differential abundance (LinDA) analysis between each supplementation group compared to control within each diet (FR, left panel; FF, middle panel; DI, right panel). Plot includes the intersection size per intersection in the top, with the indication of group or comparison at the bottom right and the set size of each group at the bottom left. orange: p-value *<* 0.1 from two-sided t-test; blue: FDR-adjusted p-value by Benjamini-Hochberg adj.p*<* 0.05. **C)** Dotplots of LinDA regression coefficients as showing supplementation-affected features, calculated based on longitudinal data from baseline (week 0), week 1 and week 2 of supplementation. Supplementation-specific trends during FF diet shown in central panel. Primary filtering for FP- associated effects included only coefficients with p-values *<* 0.1, with unidirectional signs in all FP- supplemented groups, followed by the exclusion of coefficients total amount *<* 0.5, as well as coefficients with standard errors (SE) exceeding zero. Subsequent secondary filtering performed with similar thresholds but for PP-associated effects, followed by tertiary filtering for AV-associated affects only. In a next step, unfiltered regression coefficients between FF and FR (ctrl) and DI and FF (ctrl) were added for the previously filtered taxa in the left and right panel, respectively. Error bars indicating SE, dot size reflecting statistical trends (*−log*_10_ (*P* value)), with filled dots indicating statistical significance with FDR-adjusted p-value*<*0.05. Heatmap colors indicating mean relative abundance in % of each species at baseline (wk 0) across all FF fed mice. Effect mediator color-scale indicates if the effect was FP-mediated (dots for all three groups), PP-mediated (dots for PP-supplemented groups only), or AV-mediated (single dot for AV-supplementation).

The effect of the supplements on *A. muciniphila* extended to the small intestine, where its relative abundance was significantly lower in the FP and FP+PP groups compared to the control (Supp 2D). Crucially, this effect appeared mainly FP-mediated as 1) the additional PP-supplementation showed no additional shifts, and 2) the DI group showed no significant change in *A. muciniphila* abundance in this gut region (Supp 2D).

Given the distinct physiological environment of the small intestine–characterized by rapid transit times, lower bacterial loads, higher oxygen tension, and direct exposure to host factors like bile acids and immune effectors [96], these findings support different underlying mechanism. The data imply that the changes in *A. muciniphila* may be driven by the postbiotic itself, or direct host-mediated responses to the bioactive components of the supplements, rather than solely by microbe-microbe competition for dietary fiber [97].

In addition to its effects on *A. muciniphila*, the fecal microbiota analysis revealed that FP supplementation consistently modulated two other taxa during the FF diet. First, the supplementation counteracted the FF-driven increase of *P. goldsteinii* (Figure 2C, Supp 2A), an opportunistic mucin-degrader with immunoregulatory potential [94, 95, 98, 99]. This parallel effect on two distinct mucus-degrading and utilizing bacteria suggests that the supplement may exert a broader protective effect on the mucus barrier by attenuating the expansion of such species during fiber deprivation.

Second, the FP supplementation reversed the FF-associated decrease of an unclassified *Enterocloster* species (Figure 2C, Supp 2A). This genus, part of the *Lachnospiraceae* family, includes species that produce metabolites critical for host-microbe homeostasis. For instance, members of this genus are known to produce indole-3-propionate, thereby inducing regulatory T cells, which enhances epithelial barrier function [100–105]. Therefore, by counteracting the depletion of this genus, the supplement may promote gut health through microbe-mediated support of mucosal immunity and barrier integrity.

Apart from the FP-mediated effects, our analysis identified several taxa that were consistently modulated by PP supplementation during the FF diet. The effects could be broadly categorized based on their similarity to the fiber reintroduction seen in the DI group.

A number of PP-mediated changes resembled the effect of dietary fiber reintroduction, suggesting direct fiber-induced effects. Specifically, PP reversed the FF-associated decrease of members of several genera, including *Alistipes*, *Laedolimicola*, *Oribacterium*, as well as *Mucispirillum schaedleri* (Figure 2C, Supp 2A). This aligns with literature associating *Alistipes* and *Oribacterium* with high fiber intake [106–108]. Notably, the fiber-dependent role of *M. schaedleri* in mucosal health is well known. Contrary to our results, Kuffa et al. showed how a FF exclusive enteral nutrition (EEN) limited the ability of *M. schaedleri* to occupy its niche in the mucus layer, thereby alleviating colitis in a Crohn’s like mouse model [109]. However, this effect was highly dependent on a cross-feeding mechanism with *Ruminococcus torques*—an almost undetected taxa among our samples—indicating that diet-modulated microbial shifts will always depend on specific competitive and cooperative mechanisms of the underlying community [10, 109]. PP supplementation led furthermore to a decrease in *Alloprevotella*, a response that also resembled fiber reintroduction (Figure 2C, Supp 2A), although species of this genus are commonly more associated with high-fiber diets [110, 111].

In contrast, the modulation of other taxa by PP did not align with the changes observed in the DI group, indicating responses that may be specific to certain fiber-types within the prebiotic. For instance, PP supplementation increased *Dorea* and *Cryptobacteroides* (Figure 2C, Supp 2A). The increase in *Dorea* is consistent with its known responsiveness to dietary fiber [112–115]. While we do not see an increase in mean relative abundance upon 6 weeks of fiber-reintroduction (DI) compared to the baseline (Figure 2A), there might be an underlying community-specific effect in that case. An increase in *Schaedlerella* was also noted in the longitudinal analysis, however, this was an artefact due to much higher baseline abundance in the FF placebo-intervened and FP-supplemented groups compared to the other two groups, which led to no differences in abundance in the cross-sectional data at week 2 (Figure 2C, Supp 3A, Supp 2A).

Finally, the analysis highlighted considerable changes within one characterized species of the *Alistipes* genus. The PP-supplementation robustly increased the abundance of one species (*Alistipes A 871400 sp002362235*) compared to FF, which resembled the shift during DI (Fig 2C, Supp 2A). Studies show a dietary fiber-responsiveness of this genus in humans [106, 107]. Notably, all three supplementation combinations led to an additional increase in this species in the DI compared to the placebo-intervened DI group, indicating overall product-mediated effects on this particular species (Supp 1C).

Analysis of the full supplement combination group (FP+PP+AV) revealed that the addition of AV was associated with trends of change in 15 additional taxa compared to the FF control (Figure 2C). Although these trends did not reach statistical significance after FDR correction, the majority of them, particularly decreases in certain taxa, mirrored the shifts observed upon fiber reintroduction in the DI group. This suggests that AV supplementation may induce community changes that partially overlap with those driven by dietary fiber.

To identify the most reliable signals, we focused on taxa whose trends were consistent across both the longitudinal (Figure 2C) and cross-sectional (Supp 2A) analyses. This stringent approach highlighted four taxa that showed consistent decreases: *Duncaniella muris*, *Odoribacter splanchnicus*, *JAGBWK01* spp., and *Muribaculum* spp. With the exception of *D. muris*, the observed decreases in these taxa were also consistent with the effect of fiber reintroduction in the DI group.

The observation that AV supplementation induces microbial shifts that resemble those of dietary fiber points towards a potential overlap in biological function. While aloe polysaccharides have been shown earlier to modulate the gut microbiome [21, 22], we were the first to show, that these changes partly resemble those achieved by dietary fiber intervention.

Notably, the full supplement combination also induced changes not observed in the DI group. These included consistent increases in the butyrate-producing genera *Dysosmobacter* and *Ki-neothrix* [116–118] (Figure 2C). While these taxa slipped our analysis that was designed to detect robust changes, a closer look revealed that both taxa did respond negative to fiber, and specifically showed an increase in relative abundance in the FP, which was first counteracted by the additional supplementation of fiber, and then again enforced by the AV-supplementation (Supp 2A, Supp 3B-C). Our data suggests therefore a strong negative association with fiber, that is counteracted by a strong positive responsiveness to the FP and AV supplement. Interestingly, species within these genera have been previously linked to reduced inflammation in mice, indicating their potential to contribute to gut barrier protection [116, 119]. Overall, these data indicate a potential microbiome-mediated anti-inflammatory and gut barrier protective effect of the postbiotic and AV supplement, which out compete positive effect of fiber-introduction for certain species. This hypothesis would make them candidate-supplements to be tested in the combination of FF EEN–to date one of the single effective interventions in the treatment of IBD [120, 121].

Finally, we examined the effects of supplementation in the stable, fiber-rich environment of the FR-fed mice. In this context, few consistent taxonomic shifts were observed; while some trends were associated with the FP and FP+PP+AV groups, no single association passed our stringent filtering criteria for the PP-only supplementation (Supp 2B). This lack of marked changes in a stable, fiber-replete community reinforces the conclusion that the supplements exert their most profound effects on a microbiome that has been pre-perturbed by a low-fiber diet.

Overall, our data demonstrate that a postbiotic supplementation, alone or in combination with a prebiotic and aloe vera, can drive specific microbial changes in a low-fiber perturbed gut community. Crucially, many of these changes recapitulated effects that were otherwise only achieved by the complete reintroduction of dietary fiber, and a few indicate additional positive changes, highlighting the potential of such supplements to restore key functions in a dysbiotic microbiome.

### Postbiotic-based supplementation leads to functional microbial changes with possible implications for the host

To elucidate the functional consequences of supplementation and explore the mechanisms driving the observed taxonomic shifts, we performed 16S-based transcript predictions (Supp 3D-E). The analysis suggested a functional remodeling of the microbiota that corresponded with, and likely underpinned, the changes in community structure, which we confirmed with metatranscriptomic data on the cecal contents of the FF fed mice (Figure 3, Supp 4). Notably, the transcriptomic data confirmed the lower relative abundance of key mucin-degrading bacteria, such as *A. muciniphila* and the increase of *Alistipes spp.* (Supp 2E). It further provided a mechanistic link by showing a concurrent downregulation of their mucinolytic machinery, as contributed by *A. muciniphila* and *P. goldsteinii* (Figure 3A-B, Supp 4). These alterations were part of a broader, community-wide metabolic pivot from host-glycan degradation toward the utilization of supplemented prebiotics and their fermentation products.

**Figure 3:**
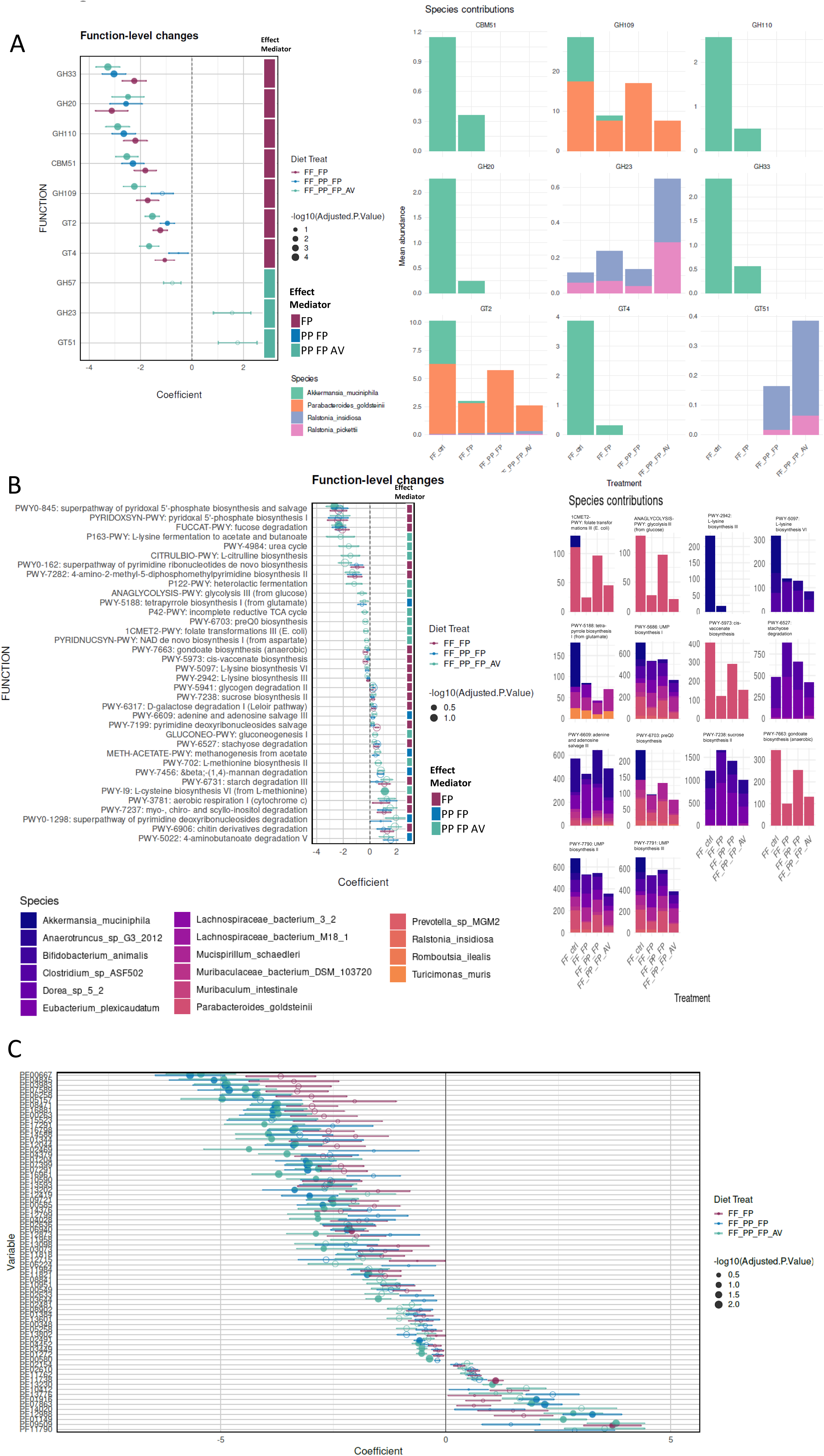
A-C) Dotplots of regression coefficients from linear models for differential abundance (LinDA) of metatrascriptomic A) CAZyme B) pathway C-D) PFAM function abundance changed over time during supplementation in the FF fed mice compared to the unsupplemented control group. Error bars indicating standard error (SE), dot size reflecting statistical significance (*−log*_10_ (*P* value)), with filled dots P*<*0.05, heatmap colors indicating mean relative abundance in % of each function at baseline (wk 0) across all FF fed mice. Functions were filtered for P*<*0.1 threshold and only shared functions between supplementation and control group were included. Species contributions as total abundance in counts per million indicated in right panel for A) and B) or in **Supplementary** Figure 4 for C). Effect mediator color-scale indicates if the effect was FP-mediated (dots for all three groups), PP-mediated (dots for PP-supplemented groups

A primary indicator of this functional shift was the decreased expression of microbial carbohydrate-active enzymes (CAZymes) associated with mucin degradation (Figure 3A). We observed a significant reduction in transcripts for sialidases (GH33), enzymes specific to mucin breakdown, which were contributed entirely by *A. muciniphila* [122]. Similarly, transcripts for alpha-N-acetylgalactosaminidases (GH109) and alpha-galactosidases (GH110), which also target host glycans, were significantly reduced and primarily expressed by *A. muciniphila* and *P. goldsteinii* (Figure 3A) [93, 122]. The expression of carbohydrate-binding module CBM51, a domain often appended to mucin-targeting enzymes and solely expressed by *A. muciniphila* in this context, was also significantly lower in supplemented groups (Figure 3A). At the pathway level, we observed a concordant decrease in the expression of the fucose degradation pathway (FUCCAT-PWY) (Figure 3B), indicating reduced breakdown of fucosylated mucin O-glycans.

These findings align with previous work showing that mucolytic activity increases under fiber-free conditions [11], an observation supported in our study by *in vitro* assays showing elevated sulfatase, *α*-N-Acetylglucosaminidase, and fucosidase activity on the FF diet (Supp 3F). However, while our transcriptomic data suggested a counteraction by the supplements, these functional shifts were not confirmed at the level of *in vitro* enzymatic activity, where no significant reductions were observed (Supp 3F). This discrepancy may be explained by several factors; notably, the activity of sialidases, which showed the most pronounced change at the transcript level, was not assayed due to substrate limitations. Furthermore, the lack of change in other enzymes, such as *α*-galactosidase, may reflect their broader substrate specificities or the shorter duration of our dietary intervention compared to previous studies [11].

Concurrently with the downregulation of mucin-degradation machinery, we observed a consistent upregulation of pathways associated with the metabolism of fiber and prebiotic components. All supplemented groups exhibited increased expression of pathways for starch (PWY-702), stachyose (PWY-6527), chitin (PWY6906), and inositol (PWY-7237) degradation (Figure 3B). The enrichment of inositol degradation is particularly noteworthy, as inositols are implicated in host metabolic and neurological homeostasis [123–127]. Groups receiving the PP also showed increased expression of the *β*-(1,4)-mannan degradation pathway (PYW-7456). Pfam protein family analysis further supported this shift, revealing increased expression of L-arabinose isomerase domains (PF11762, PF02610), primarily contributed by the fiber-responsive butyrate producer *Eubacterium plexicaudatum* [128, 129] (Figure 3C, Supp 4).

Beyond metabolism, supplementation was associated with a reduction in transcripts linked to microbial virulence and host inflammation. We noted decreased expression of several type II secretion system proteins (PF00263, PF05157), oxidative stress response proteins (PF07399, PF14376, PF13098), and adhesion molecules (PF02469, PF16961), with *A. muciniphila* being a major contributor to several of these changes (Figure 3C, Supp 4). Furthermore, we observed a significant reduction in pathways for L-lysine biosynthesis (PWY-2942, PWY-5097) and its fermentation to acetate and butanoate (P163-PWY), again largely driven by the decreased abundance and activity of *A. muciniphila* (Figure 3C, Supp 4). This is relevant as it suggests that changes in function of single microbes on entire community functions, which has in particular been shown for *A. muciniphila* [130]. Furthermore, Lysine has been shown to activate mTORC1, which can drive type III immune responses, thereby improving immunotolerance [131]. However, type III immune response activation can also have detrimental effects in certain conditions, as it has been shown to exacerbate colitis in a susceptible host, such as IL10-/- mice [14].

In summary, the metatranscriptomic data demonstrate that the supplements induce a profound functional remodeling of the gut microbiota under fiber-free conditions. The consistent effects observed across all three supplementation arms suggest that the postbiotic product is a key driver of these changes. The microbiome appears to shift from a reliance on host-derived mucins to the utilization of externally provided substrates, a change accompanied by a downregulation of microbial functions associated with inflammation. These findings once again indicate that postbiotic-based supplements can recapitulate key functional benefits of dietary fiber, offering a promising strategy for microbiome modulation in clinical settings where high-fiber diets are contraindicated, such as for patients receiving EEN [120].

### Functional community shifts show potential to counteract barrier integrity impairment and diet-induced type III immune response cytokine expression in fiber deprived mice

Finally, we investigated whether the observed microbial functional shifts translated to changes in host physiology. Histological analysis of the distal colon revealed that mice receiving the full supplement combination (FP+PP+AV) exhibited a trend towards a thicker mucus layer compared to the FF controls (One-way ANOVA of supplementations vs. control, without correction for multiple comparison (Fisher’s LSD test) Figure 4A). This anatomical observation is consistent with our molecular data showing a supplement-driven reduction in the abundance of mucin-degrading bacteria and the expression of microbial mucinolytic enzymes, supporting a direct link between the supplement’s modulation of microbiota function and the preservation of this critical barrier component [13]. While we detected no significant differences in the expression of key barrier integrity markers (e.g., Occludin (Ocln), trefoil factor family peptide 3 (Tff3), Krüppel-like factor 4 (Klf4)) across the FF groups, the upregulation of repair-associated factors such as fucosyltransferase 2 (Fut2), Tff3, and Klf4 in single control animals may reflect an early compensatory response to barrier stress, a response that was less apparent in supplemented mice.

**Figure 4:**
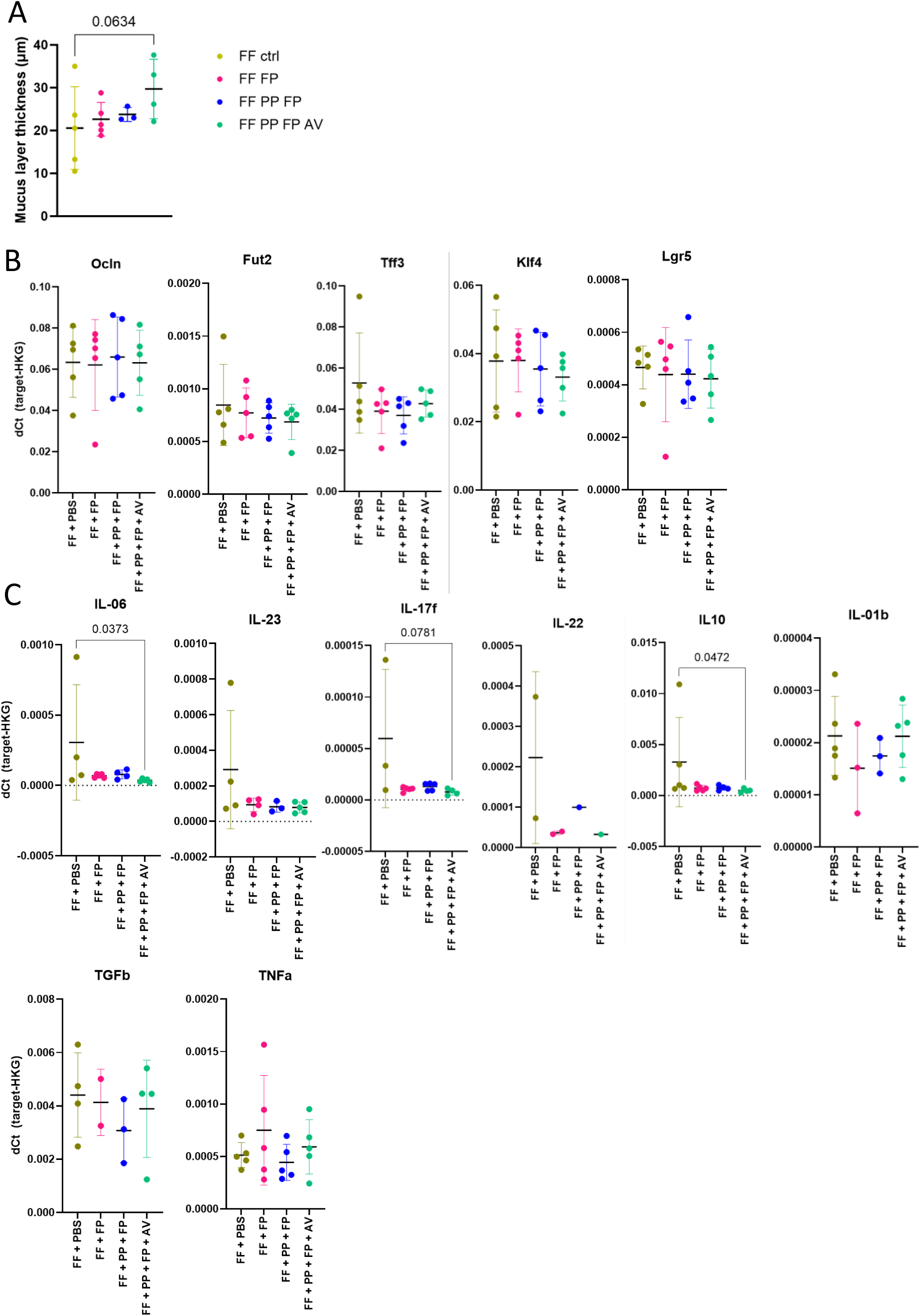
A) Individual dotplot of average mucus layer thickness in µm per mouse fed the FF diet with mean indicated as horizontal bar and standard deviation indicated as error bars. Statistical trend determined by one-way ANOVA of supplemented vs. control groups, without correction for multiple comparison (Fisher’s LSD test, p*<*0.1) **B-C)** Individual dot plots of efficiency-corrected delta Ct values of target **B)** membrane integrity marker and **C)** immuno cytokines against Gapdh housekeeping gene. Ct values exceeding the reliable detection threshold identified by target amplification of sample pre-dilutions were excluded from the analysis (Ocln*>*30.5, Fut2*>*31.5, Tff3*>*29, Klf4*>*32, Lgr5*>*32.5, IL-6 Ct*>*35, IL-23 Ct*>*33, IL-17f Ct*>* 34, IL-22 Ct*>*33, IL-10 Ct*>*35, TNFa Ct*>*31, IL-12 Ct*>*35, TGFb Ct*>*33). Statistical analysis was performed for all n*>*3 comparing supplemented group vs. control using a one-way ANOVA for normal distributed values and Kruskal-Wallis nonparametric test with for abnormal distributed values (IL-6, IL-23, IL-12, IL-17f, IL-10) with post-hoc multiple comparison correction by Benjamini-Hochberg (Q *<* 0.05).

This preservation of barrier integrity was accompanied by a notable attenuation of host immune activation. In stark contrast to FR-fed mice, whose colonic cytokine expression remained below detection limits, FF-fed control mice exhibited an upregulation of type III immune response cytokines (Interleukin-6 (IL-6), IL-23, IL-17f, IL-22) and the immunoregulatory cytokine IL-10. Supplementation significantly dampened this response, with the full supplement group showing significantly lower expression of IL-6 and IL-10 compared to controls. This attenuated immune phenotype is consistent with our metatranscriptomic findings of reduced microbial L-lysine metabolism, a potential upstream modulator of type III immunity [131]. The simultaneous elevation of pro-inflammatory and regulatory cytokines in the FF control group suggests that the fiber-deprived gut environment provides a potent stimulus that necessitates a counter-regulatory response to maintain homeostasis. We strongly emphasize on the fact, that our observations were done in a healthy mouse model, that cannot predict pathogenic mechanisms. In a genetically susceptible host, such as an IL-10-deficient model, this balance may fail, leading to overt pathology [132]. Overall, the reduced marker gene expression in supplemented mice compared to the controls may reflect an early preservation of gut barrier integrity under fiber-deprived conditions, potentially limiting the need for compensatory host-driven immune or repair responses.

## Conclusion

In conclusion, our study demonstrates that a postbiotic-based supplementation can effectively steer the composition and function of a gut microbial community destabilized by fiber deprivation. The intervention drove a significant metabolic shift within the microbiome, away from the degradation of host mucus glycans and towards the utilization of dietary prebiotics and supplemented postbiotics. This functional remodeling, highlighted by the counteracted increase of mucolytic bacteria like *A. muciniphila*, was associated with the preservation of the colonic mucus layer and a marked reduction in the low-grade, type III immune activation characteristic of a fiber-free diet. These findings underscore the potential for microbial fermentation products to directly modulate host-microbe interactions, thereby recapitulating some of the key benefits of dietary fiber [97]. While our results present a promising strategy for supporting gut health, the precise outcomes of such interventions are likely to be context-dependent, shaped by the initial state of an individual’s microbiome and host-specific factors [133]. Nevertheless, this work supports the development of postbiotic-based supplements as a means to promote intestinal homeostasis, particularly in contexts where dietary fiber intake is limited.

## Acknowledgments

We gratefully acknowledge the funding of this research by MEDICE Arzneimittel Pütter GmbH & Co. KG., and the support of the Luxembourg National Research Fund (FNR) through CORE grants (C15/BM/10318186 and C18/BM/12585940) and BRIDGES grant (22/17426243) to M.S.D. We also thank Nathalie Nicot at the LuxGen Platform of the Luxembourg Institute of Health and the Laboratoire National de Santé for her support in the metatranscriptomic sequencing, Alexander Hundt and his team at the IBBL for processing of histology slides, and Anaïs Oudin and her animal facility team of LIH, especially Marion da Costa, for their technical support during the mouse experiment. For the purpose of open access, and in fulfillment of the obligations arising from the grant agreement, the author has applied a Creative Commons Attribution 4.0 International (CC BY 4.0) license to any Author Accepted Manuscript version arising from this submission.

## Declarations

### 0.1.1 Availability of data and materials

The raw files from full-length 16S rRNA gene sequencing and RNA sequencing have been deposited in the European Nucleotide Archive (ENA) at EMBL-EBI under accession number PRJEB97506 (https://www.ebi.ac.uk/ena/browser/view/PRJEB97506).

Supplementary data tables are available for download: 10.5281/zenodo.17182513

### 0.1.2 Disclosure statement

M.S.D. works as a consultant and an advisory board member at MEDICE Arzneimittel Pütter GmbH & Co. KG. U.B. is managing director and R.A. managing partner at MEDICE Arzneimittel Pütter GmbH & Co. KG.

### 0.1.3 ORCID

Mahesh S. Desai http://orcid.org/0000-0002-9223-2209

### 0.1.4 Authors’ contributions

Conceptualization, A.R., E.T.G, A.S., A.B., R.A., U.B., and M.S.D; Data curation, A.R., E.T.G, and O.H.; Formal analysis, A.R.; Experiments, A.R., S.W., and A.D.; Investigation, A.R., E.T.G, O.H., S.W., A.D., C.D., A.B., R.A., U.B., and M.S.D; Methodology, A.R., E.T.G, S.W., A.D., O.H. and C.D.; Resources, U.B., and M.S.D.; Writing – Original Draft, A.R.; Writing – Review Editing, A.R., E.T.G, O.H., S.W., A.D., C.D., A.B., R.A., U.B., and M.S.D; Supervision, M.S.D.; Funding Acquisition, R.A., U.B., and M.S.D.

---

**Supplementary Figure 1:**
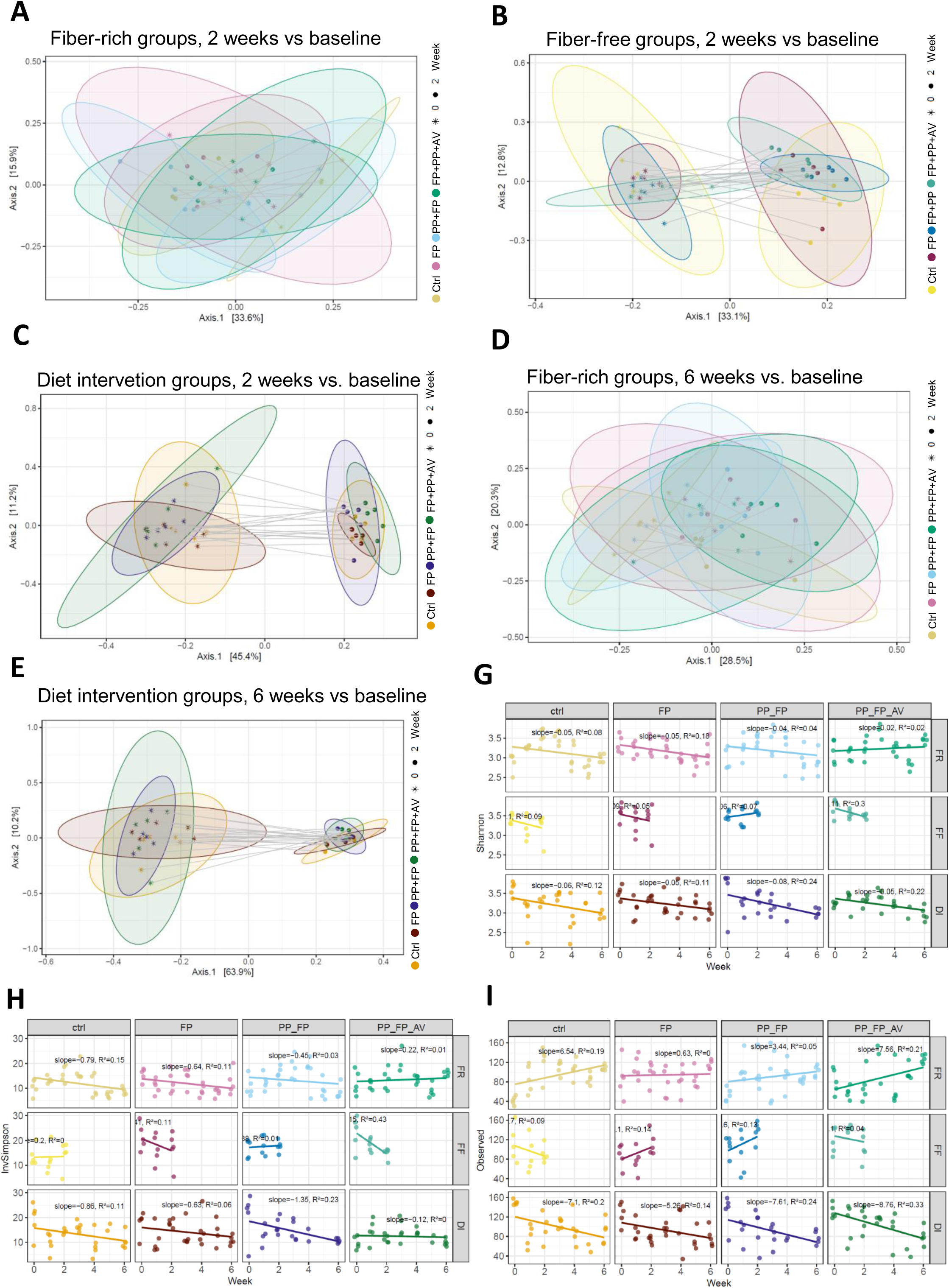
**A-E)** Principal coordinate analysis (PCoA) ordination of a Bray-Curtis dissimilarity matrix based on the relative abundances of fecal samples per mouse per week at baseline (wk 0) and after 2 weeks A) fiber-rich (FR), B) fiber-free (FF) diet or C) dietary fiber intervention (DI), or after 6 weeks D) FR or E) DI with each n=20 mice per diet per time point, resulting in n=40 per plot. **G-I)** Individual dotplots of alpha-diversity measures (G) Shannon, H) Inversed Simpson, I) Observed) for each mouse at baselines and weekly during supplementation with FP, FP+PP and FP+PP+AV or placebo-intervention (ctrl), with correlation analysis over time, slopes and R2 indicated.

**Supplementary Figure 2:**
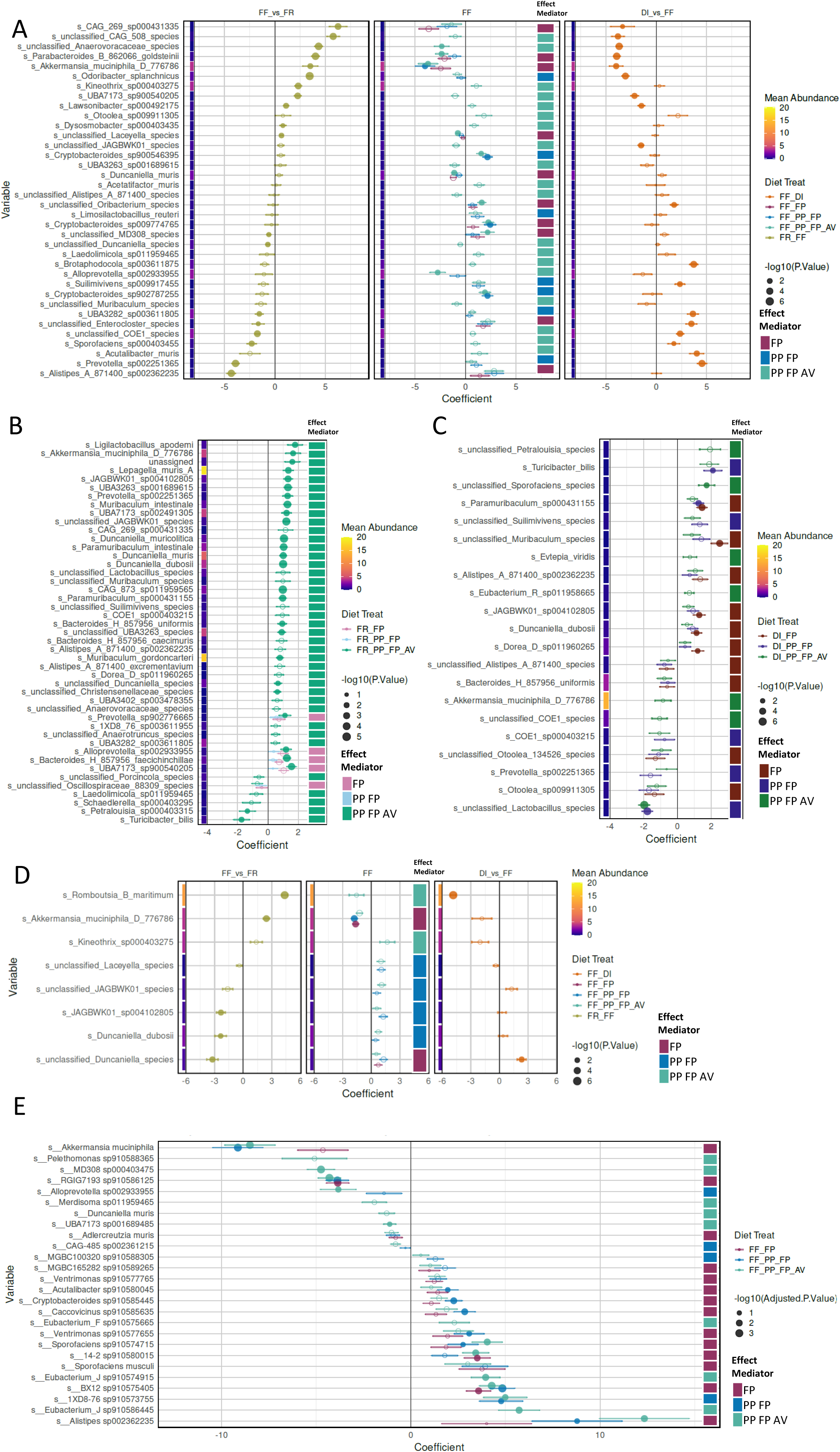
**A-E)** Dotplots of linear models for differential abundance (LinDA) regression coefficients showing supplementation-affected features based on A, D, E) cross-sectional data at week 2, or longitudinal data from baseline (week 0), week 1 and week 2 (B-C). Supplementation-specific trends during A) fiber-free diet in the central panel compared to diet-dependend trends in the left (fiber-rich (FR) vs. FF) and right (diet-intervention (DI) vs. FF) panel; B) during FR diet, and C) during DI. Analysis based on compositional 16S rRNA sequencing data of A-C) fecal samples, D) small intestinal contents, and E) metatranscriptomic data of cecal contents. Primary filtering for FP-associated effects included only coefficients with p-values *<* 0.1, with unidirectional signs in all FP-supplemented groups, followed by the exclusion of coefficients total amount *<* 0.5, as well as coefficients with standard errors (SE) exceeding zero. Subsequent secondary filtering performed with similar thresholds but for PP-associated effects, followed by tertiary filtering for AV-associated affects only. Error bars indicating SE, dot size reflecting statistical trends (*−log*_10_ (*P* value)), with filled dots indicating statistical significance with FDR-adjusted p-value*<*0.05. A-C) Heatmap colors indicating mean relative abundance in % of each species at baseline (wk 0) or at week 2 (D) across all FF fed mice. Effect mediator color-scale indicates if the effect was FP-mediated (dots for all three groups), PP-mediated (dots for PP-supplemented groups

**Supplementary Figure 3:**
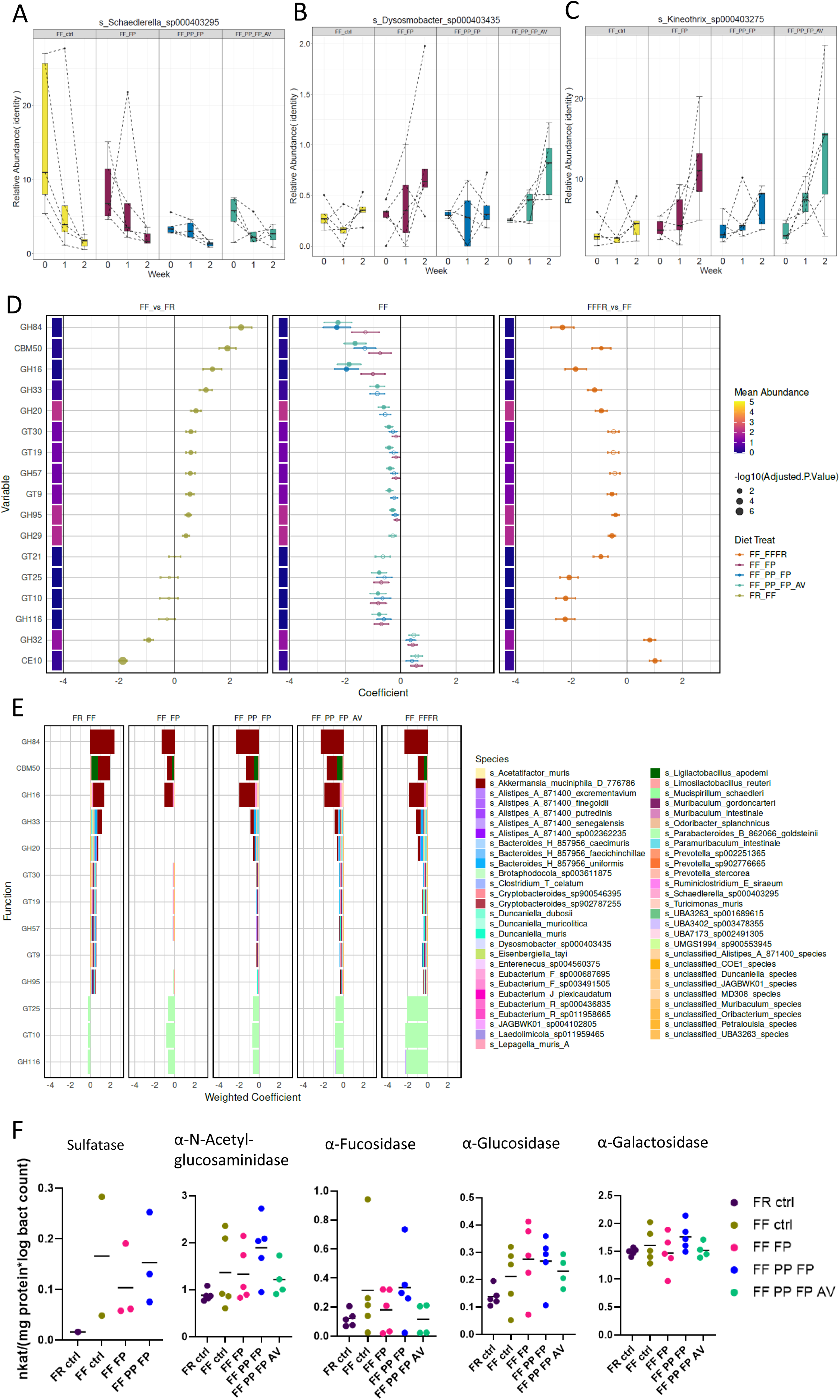
**A-C)** Boxplots of group relative abundances of group-specific species changes over time during fiber-free (FF) diet supplementation with a fermented postbiotic (FP; FF FP), together with a prebiotic (PP; FF PP FP), and additional aloe vera (AV; FF PP FP AV) compared to PBS (ctrl) over 1 and 2 weeks compared to baseline (0 weeks) with connecting lines indicating single mice relative abundances. A) For *Schaedlerella spp.*, B) for *Dysosmobacter spp.*, C) for *Kineothrix spp.*. **D)** Dotplots of regression coefficients from linear models for differential abundance (LinDA) of 16S rRNA sequencing based CAZyme transcript predictions in the FF fed mice during supplementation with FP, PP and AV (central panel), or compared to a fiber-rich (FR) diet (left panel) or diet intervention (DI) (right panel). Error bars indicating standard error (SE), dot size reflecting statistical significance (*−log*_10_ (FDR-adjusted*P* value)), with filled dots P*<*0.05, heatmap colors indicating mean relative abundance in % of each function at baseline (wk 0) across all FF fed mice. Functions were filtered for P*<*0.1 threshold and only shared functions between supplementation and control group were included. Effect mediator color-scale indicates if the effect was FP-mediated (dots for all three groups), PP-mediated (dots for PP- supplemented groups only), or AV-mediated (single dot for AV-supplementation). Species contributions indicated by stacked barplots in **E)**. **F)** Individual dotplots of enzyme activities per mg protein, normalized by the log bacterial count. n=5 per group, except for FP+PP+AV n=4, because of insufficient amount of fecal samples. Data excluded for Sulfatase in case of insufficient protein concentration.

**Supplementary Figure 4:**
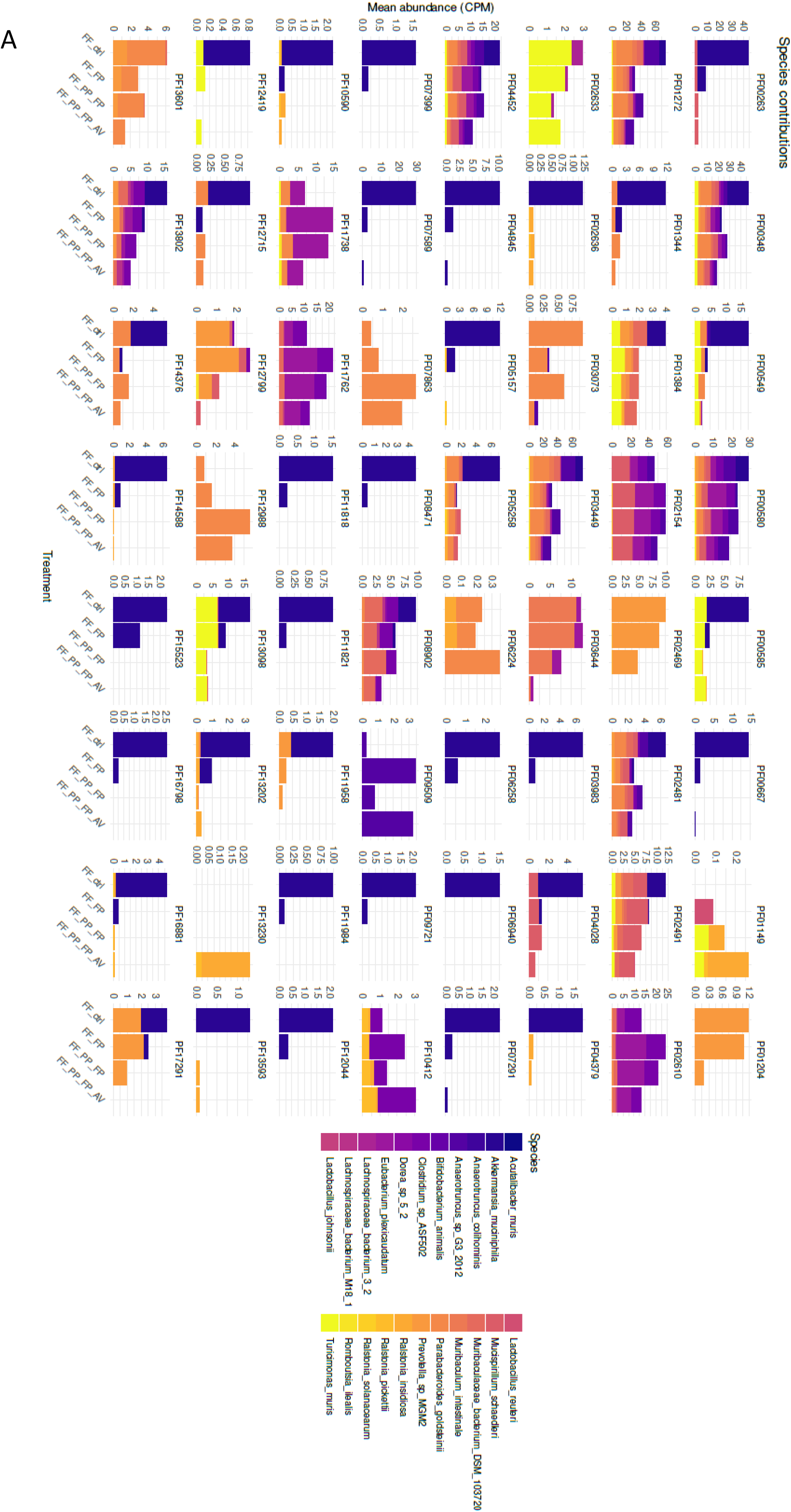
Stacked barplots of total abundance in counts per million of species contributing to changed Pfam transcripts in Figure 3 **C)** from linear models for differential abundance (LinDA) of metatrascriptomic cecal contents in the FF supplemented fed mice compared to the un- supplemented control group.

## Notes

### Competing Interest Statement

M.S.D. works as a consultant and an advisory board member at MEDICE Arzneimittel Puetter GmbH & Co. KG, Germany. A.B. is employed by MEDICE, U.B. is managing director and R.A. is a managing partner at MEDICE Arzneimittel Puetter GmbH & Co. KG.

